# Targeted degradation of *Drosophila* FOG homolog U-shaped mimics macrophage transdifferentiation in S2 cells

**DOI:** 10.1101/2025.05.19.654861

**Authors:** Deborah Trummel, Jonathan Lenz, Wiebke Milani, Lara Sophie Joost, Lotta Nele tom Dieck, Sven Bogdan, Alexander Brehm

## Abstract

The highly plastic cellular component of the *Drosophila* immune system consists of three main blood cell types - plasmatocytes, crystal cells and lamellocytes - that together allow effective responses to various insults. Infection with parasitic wasp eggs results in a rapid increase in highly specialized lamellocytes that are generated by differentiation from hemocyte precursors as well as by transdifferentiation from plasmatocytes. How differentiation and transdifferentiation are regulated at the molecular level is not well understood. Here, we show that inducible degradation of the Friend of GATA (FOG) homolog U-shaped (Ush) in the plasmatocyte-like S2 cell line results in downregulation of plasmatocyte marker genes and subsequent upregulation of lamellocyte marker genes. This transcriptional shift is accompanied by morphological and functional changes consistent with lamellocyte cell identity, including increased cell spreading and adhesion, driven by enhanced integrin expression associated with increased focal adhesions. Our findings demonstrate that targeted Ush depletion is sufficient to reprogramme plasmatocyte-like S2 cells toward a lamellocyte-like state, thereby providing the first *in vitro* model for studying processes involved in macrophage transdifferentiation.

## Introduction

The hematopoietic system of *Drosophila* produces three main blood cell (or “hemocyte”) types that originate from undifferentiated hemocyte progenitors: Plasmatocytes, lamellocytes and crystal cells [1–3]. Recent single cell transcriptomics analyses have revealed that each of these three hemocyte classes is heterogeneous and encompasses several distinct subpopulations [4–8]. Plasmatocytes show a range of activities shared by mammalian macrophages. These include removal of bacteria and cellular debris by phagocytosis, remodeling of tissues during metamorphosis and participation in wound healing. Lamellocytes combat larger parasites by attaching to their surface and by forming a capsule around them. Crystal cells mediate melanization that contributes to encapsulation. Healthy animals contain mostly plasmatocytes, a small percentage of crystal cells but almost no lamellocytes. Lamellocyte differentiation is induced by acute insults such as infection by parasitic wasps. Remarkably, lamellocytes are generated in large numbers within hours of infection by differentiation from hemocyte precursors in the lymph gland as well as by transdifferentiation from plasmatocytes [9,10]. This involves dramatic changes to cell morphology and function. Lamellocytes flatten, increase in size and acquire integrin-mediated adhesive properties that allow stable attachment to wasp eggs [11]. The molecular basis of these rapid differentiation and transdifferentiation processes is not understood owing in part to the small number of lamellocytes available for analysis and the lack of a suitable cell culture model. Unlike lamellocytes and crystal cells, plasmatocytes can be mitotically active and, thus, their propagation in the animal does not exclusively rely on undifferentiated hemocyte progenitors [3]. Moreover, cell lines derived from hemocyte precursors are similar to plasmatocytes based on their gene expression profiles and properties [12,13]. In particular, two well-established embryonic cell lines, S2 and Kc cells divide rapidly, express plasmatocyte marker genes and are characterized by robust phagocytosis activity [14–16]. By contrast, cell lines representing crystal cells or lamellocytes do not exist.

Like its mammalian homolog Friend of GATA-1 (FOG1), *Drosophila* U-shaped (Ush) is a transcriptional regulator that cooperates with GATA transcription factors and chromatin regulators during hematopoiesis [2,17]. Ush is highly expressed in hemocyte precursors and plasmatocytes. By contrast, much lower Ush levels have been reported in larval lamellocytes [18]. Moreover, loss of Ush function results in increased numbers of crystal cells in embryos and causes the spontaneous differentiation of lamellocytes in healthy larvae [19]. These findings support the idea that Ush suppresses hemocyte differentiation and that reduction of Ush expression may be a pre-requisite for differentiation and transdifferentiation.

We have recently demonstrated that Ush is expressed in S2 cells and functions as an important regulator of the S2 cell transcriptome [17]. Ush cooperates with the nucleosome remodeler dMi-2 to regulate different classes of genes including genes encoding proteins with immune function. Importantly, Ush represses key marker genes of crystal cells (*lozenge*) and lamellocytes (*atilla*) suggesting that it might suppress non-plasmatocyte transcription programmes and maintain a plasmatocyte-like cell identity. It is not known if and to what extent reduction of Ush expression could reprogramme S2 cells into the other hemocyte lineages.

Here, we established an S2 cell culture model that allows an inducible degradation of Ush protein. Cell growth assays demonstrated that degradation of Ush leads to a swift decrease in proliferation. We determined gene expression changes at several time points after induction of Ush degradation by RNA-Seq. Gene set enrichment analysis revealed that plasmatocyte marker genes are rapidly and persistently downregulated. This downregulation is followed by increased expression of lamellocyte marker genes. Moreover, microscopic analysis identified that S2 cells undergo morphological changes upon Ush degradation that are typical for lamellocytes: They become more adhesive and spread out into irregular cell shapes. RT-qPCR and immunofluorescence analysis further identified changes in integrin expression including the upregulation of lamellocyte-expressed integrins.

Taken together, we demonstrate that the targeted degradation of the hemocyte regulator Ush induces changes in plasmatocyte-like S2 cells that endow them with lamellocyte-like properties, thereby, mimicking aspects of transdifferentiation. This new S2 cell culture model will facilitate the dissection of the molecular processes underlying transdifferentiation.

## Results

### Inducible Ush protein depletion in S2 cells

We utilized a S2 cell line stably expressing GFP-tagged Ush from its endogenous locus (S2 Ush-GFP) to generate an inducible Ush degradation system [17]. Into this cell line we introduced a transgene encoding a “Degrader” protein consisting of three moieties: (1) a fragment that binds GFP with high affinity (GFP binder), (2) a F-Box protein that mediates interaction with a ubiquitin ligase complex and (3) a FKBP variant that causes ubiquitination and efficient protease-mediated degradation of the Degrader protein itself ([20], Figure 1A).

**Fig. 1:**
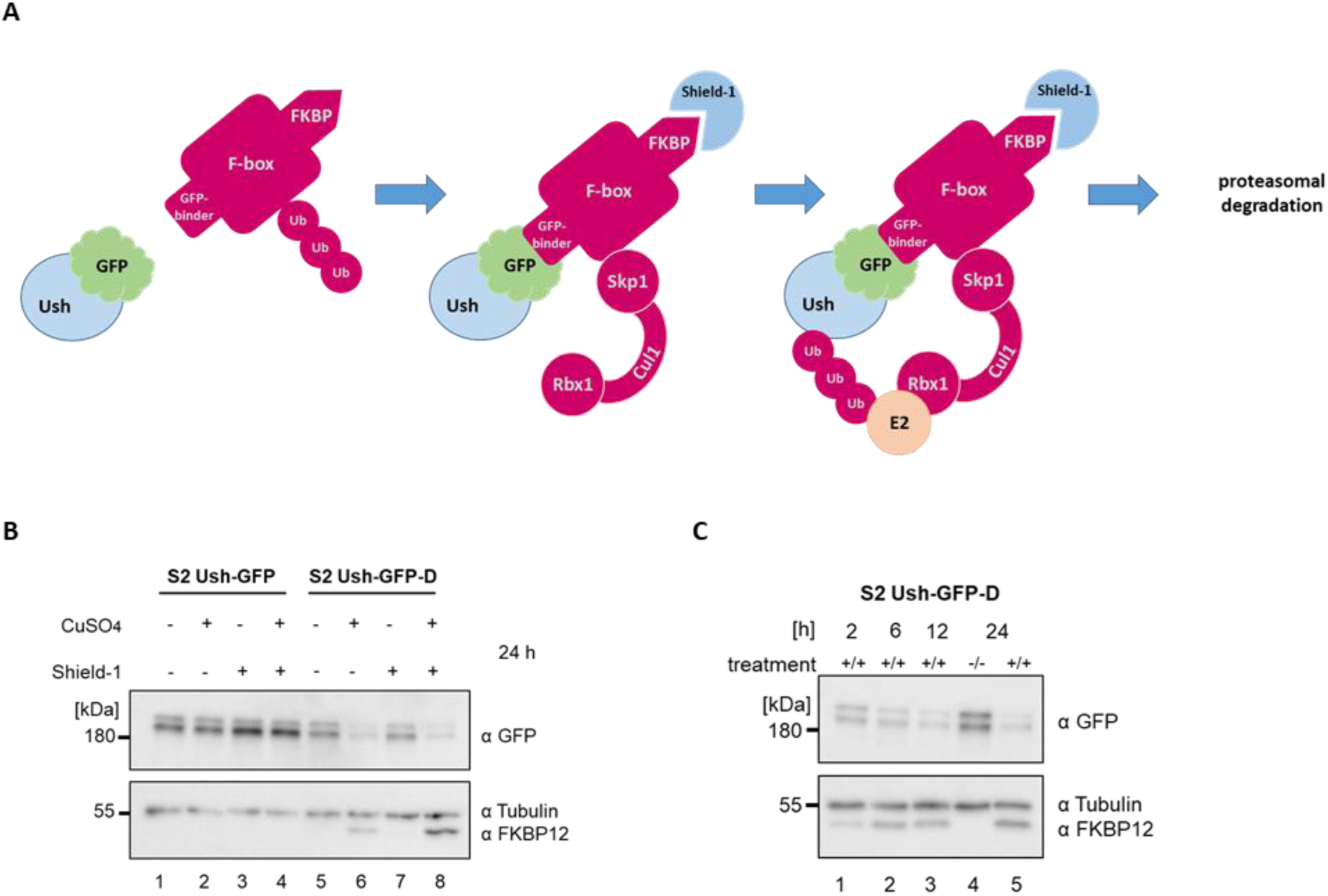
Induced degradation of Ush-GFP. **A.** The Degrader system enables depletion of GFP-tagged Ush. The FKBP-F-box-GFP binding protein fusion (Degrader, red) contains a destabilizing domain (FKBP-L106P) that results in constant degradation of the Degrader. Shield-1 (blue) inhibits FKBP and stabilizes the Degrader. This then binds GFP-tagged Ush resulting in recruitment of E2-ubiquitin-conjugating enzyme and degradation of GFP-tagged Ush. **B.** S2 Ush-GFP (parental cell line) and S2 Ush-GFP-D (Degrader expressing cell line) cells were treated with 100 µM CuSO_4_ and 5 µM Shield-1 as indicated. Whole cell lysates were prepared after 24 h and analyzed by Western Blot using the antibodies indicated on the right. Tubulin served as loading control. **C.** S2 Ush-GFP-D cells were treated with 100 µM CuSO_4_ and 5 µM Shield-1 as indicated (treatment). Whole cell lysates were prepared after the indicated time points and analyzed by Western Blot using the antibodies indicated on the right. Tubulin served as loading control.

We refer to this cell line as S2 Ush-GFP-D. In the absence of compounds that increase its expression and stability the Degrader protein was not detectable by Western blot in whole cell extracts (Figure 1B, lower panel, lane 5). Transcription of Degrader mRNA is under control of a Cu^2+^-inducible metallothionine A (MtnA) promoter [20]. Cu^2+^ induction resulted in elevated levels of Degrader that could be detected by Western Blot (Figure 1B, lower panel, compare lanes 5 and 6). Degrader concentrations were further increased by addition of Shield-1, a compound that binds FKBP and attenuates its destabilizing activity [21], Figure 1B, lower panel, compare lanes 6 and 8). Degrader binds and directs the poly-ubiquitination and subsequent proteasome-mediated degradation of GFP fusion proteins. Indeed, induction of Degrader reduced Ush-GFP levels within 24 hours (Figure 1B, upper panel, lanes 5 to 8). By contrast, treatment of control cells not expressing Degrader (S2 Ush-GFP) did not lower Ush-GFP levels (Figure 1B, upper panel, lanes 1 to 4). Expression of Degrader and concomitant reduction of Ush-GFP protein levels could be detected as soon as 2 hours after induction (Figure 1C, compare lanes 1 and 4). Degrader expression and Ush depletion increased until 24 hours after induction, but residual Ush-GFP remained detectable (Figure 1C, upper panel).

Previously, we have treated S2 cells for 96 hours with double-stranded RNA directed against Ush mRNA to deplete Ush by RNAi [17]. This approach identified different classes of genes that were activated or repressed by Ush such as genes relevant for macrophage immune function, genes encoding metabolic enzymes and genes regulating the cell cycle. Two such genes are *atilla*, which encodes a signature lamellocyte marker, and the *fatty acid 2-hydroxylase* (*fa2h*) gene, which encodes a metabolic enzyme involved in fatty acid degradation. We monitored mRNA expression of these two genes by RT-qPCR at 6, 24, 48 and 72 hours after induction of Ush-GFP protein degradation. Expression of *fa2h* was not significantly altered at the 6 hour time point but increased 5-fold after 24 hours and remained at an elevated level (Figure 2A).

**Fig. 2:**
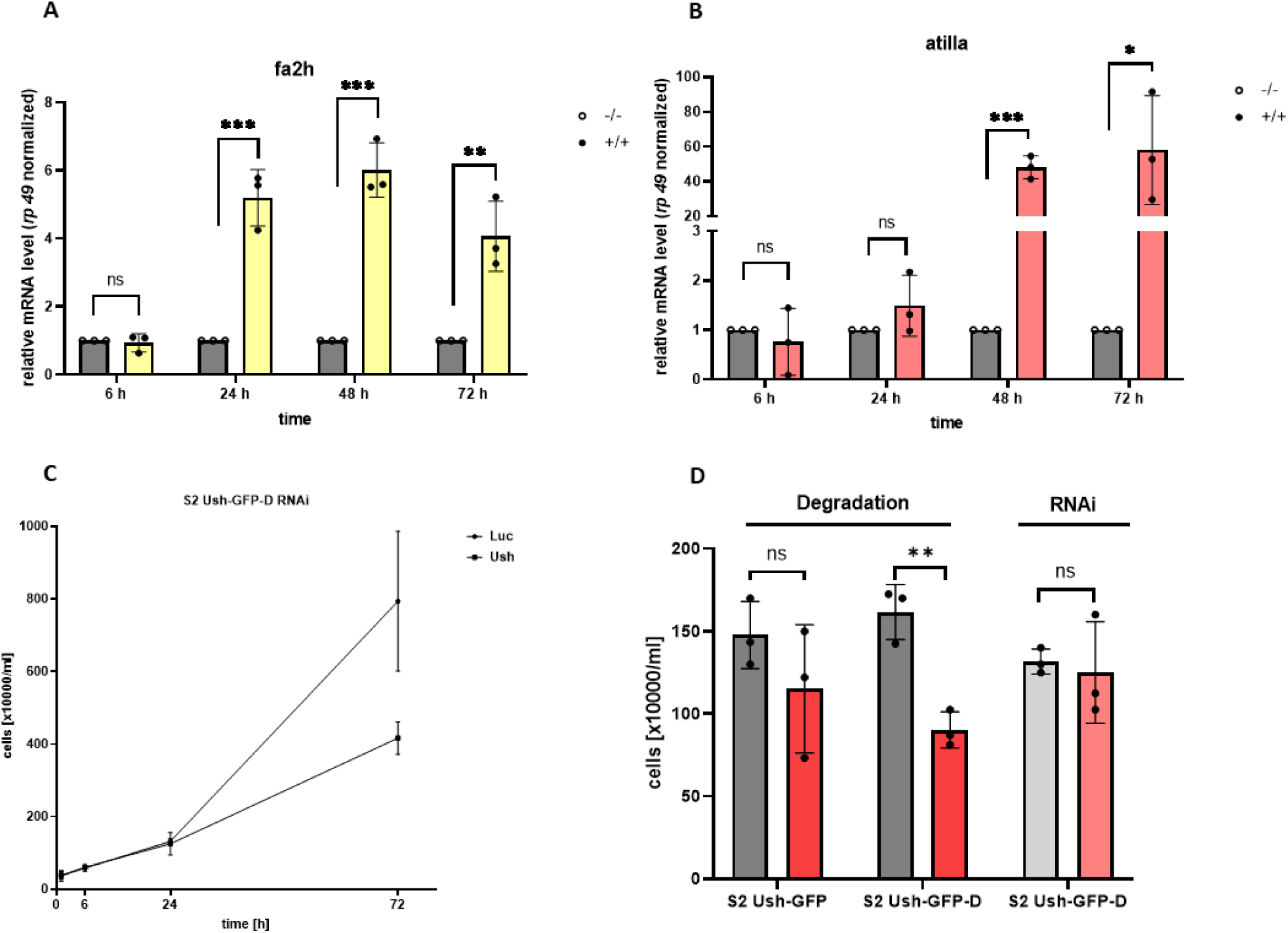
Ush-GFP degradation impairs growth of S2 cells and derepresses of Ush target genes. RT-qPCR was used to analyze mRNA levels of *fa2h* (**A**) and *atilla* (**B**) in S2 Ush-GFP-D cells following induction of Ush degradation (5 µM Shield-1, 100 µM CuSO_4_) at the time points indicated below the graph. mRNA levels were normalized to *rp49* mRNA levels. mRNA levels in untreated cells were set to 1 (grey bars). mRNA levels in treated cells (yellow and red bars) were expressed relative to these values. Individual data points and standard deviations are shown. An unpaired t-test was used for statistical analysis (ns = not significant, * p < 0.05, ** p < 0.01, *** p < 0.005). **C.** S2 Ush-GFP-D cells were subjected to Ush and Luc (control) RNAi, respectively. Cell numbers were determined at the time points indicated below the graph and plotted on the Y-axis. **D.** S2 Ush-GFP (control) and S2 Ush-GFP-D cells were treated with 5 µM Shield-1 and 100 µM CuSO_4_ (dark red bars) or left untreated (dark grey bars). Cell numbers were determined after 24 hours. Cell numbers for the 24 h time point of the RNAi experiment (C) are included for comparison (light grey and light red bars). Individual data points and standard deviations are shown. An unpaired t-test was used for statistical analysis (ns = not significant, * p < 0.05, ** p < 0.01, *** p < 0.005).

*atilla* mRNA expression was not significantly changed at both the 6 and the 24 hour time points but dramatically increased (40-fold) after 48 hours and 72 hours. This result confirms *atilla* and *fa2h* as Ush-repressed genes and demonstrates that our inducible Ush-GFP protein degradation system can resolve kinetic differences of gene deregulation.

### Ush degradation attenuates S2 cell proliferation

RNAi-mediated depletion of Ush attenuates S2 cell proliferation [17]. This effect becomes detectable 48 hours after treatment with double stranded RNA [17]. When we used RNAi to deplete Ush-GFP in S2 Ush-GFP-D cells we obtained similar results (Figure 2C). We found no significant differences in cell number when we compared cells treated with double stranded RNA directed against a luciferase control and cells treated with double stranded RNA directed against Ush-GFP at the 24 hour time point. By contrast, 72 hours after treatment cell numbers in the Ush depleted samples were reduced two-fold.

We then monitored cell numbers after induction of Ush-GFP degradation in S2 Ush-GFP-D cells. Numbers of induced cells were reduced two-fold compared to uninduced cells already after 24 hours (Figure 2D). By contrast, at the same time point cell numbers of RNAi-treated S2 Ush-GFP-D cells and copper/Shield-1-treated S2 Ush-GFP control cells were not significantly changed.

We conclude that effects of Ush depletion on proliferation can be recapitulated with our inducible Ush-GFP protein degradation system, and that the proliferation defect manifests itself more rapidly.

### Ush protein degradation alters expression of plasmatocyte and lamellocyte marker genes

Recently, several scRNA-Seq studies have determined gene expression patterns for different hemocyte populations including plasmatocyte, crystal cell and lamellocyte populations [4–8]. Hultmark and Ando have used these data to compile lists of marker genes for plasmatocytes, crystal cell and lamellocytes which show enhanced expression in the corresponding cell type [3]. We used these marker genes to ask if the plasmatocyte-like transcription programme of S2 cells is altered. We determined the transcriptome of S2 Ush-GFP-D cells 6, 24, 48 and 72 hours after induction of Ush-GFP degradation by RNA-Seq. We then performed Gene Set Enrichment Analyses (GSEA) for plasmatocyte, crystal cell and lamellocyte marker genes (Supplementary figure 1). We did not find statistically significant changes in the expression of crystal cell marker genes (Supplementary Figure 1C). By contrast, GSEA revealed a significant downregulation of the plasmatocyte marker gene set as soon as 24 hours after induction of Ush-GFP degradation (Supplementary Figure 1A). This downregulation was maintained at later time points. By contrast, lamellocyte marker genes showed an increased expression at the 24 hour time point but these changes did not cross the significance threshold (p <= 0.05; Figure 3B, Supplementary Figure 1B).

**Fig.3:**
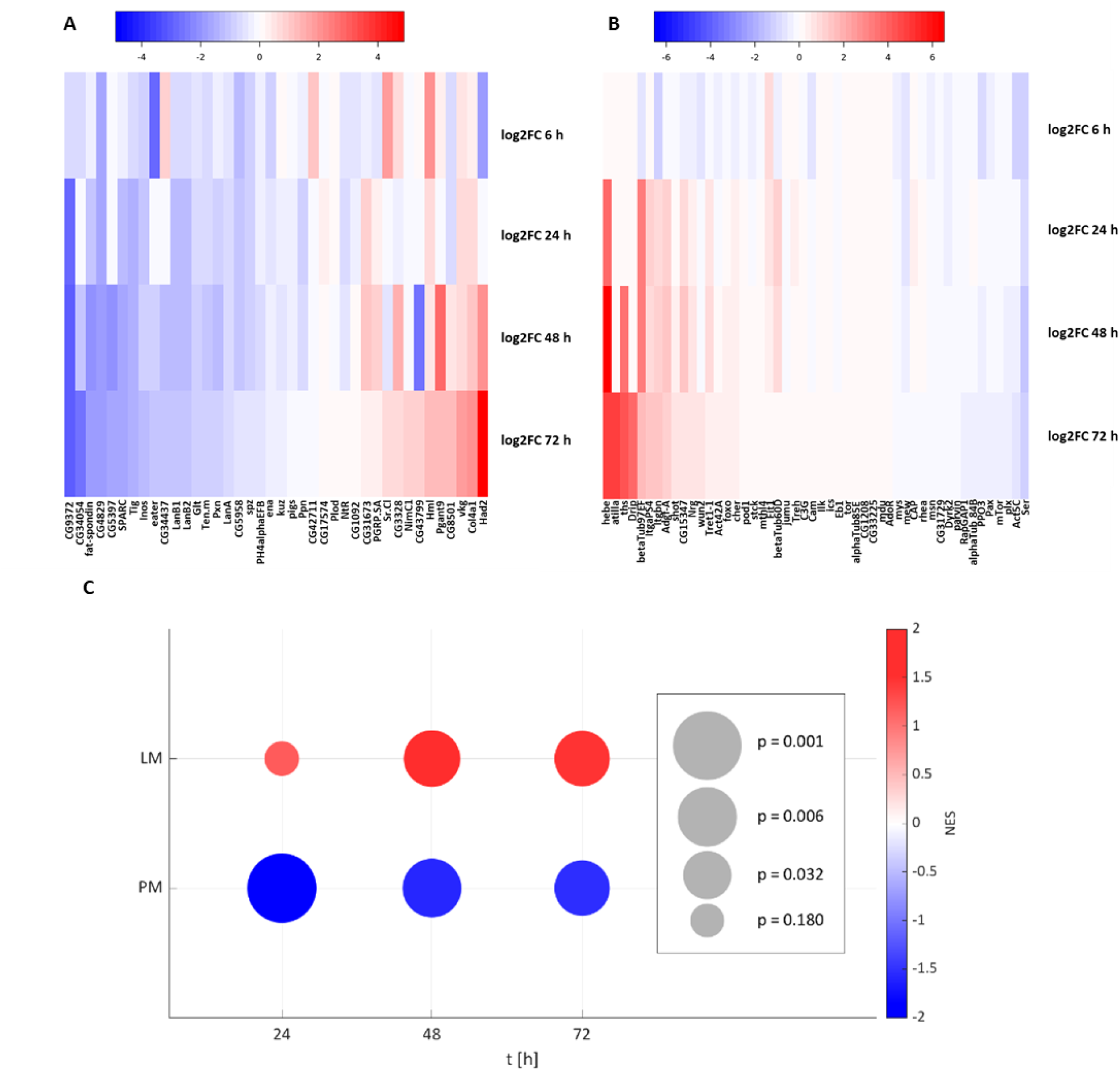
Degradation of Ush deregulates lamellocyte and plasmatocyte marker genes. S2 Ush-GFP (n=3) and S2 Ush-GFP-D (n=3) cells were treated with 5 µM Shield-1 and 100 µM CuSO_4_ to induce Ush-GFP degradation or left untreated. RNA-Seq was used to determine mRNA expression differences 6, 24, 48 and 72 h after induction. Log2-fold changes of plasmatocyte (**A**) and lamellocyte (**B**) marker genes were calculated and visualized by heatmaps. **C.** Bubble plot visualising NES (colour) and p-values (size) of GSEA (Gene Set Enrichment Analysis) data (see Supplementary figure 2 A and B, LM = Lamellocyte marker genes, PM = Plasmatocyte marker genes).

However, significant upregulation of lamellocyte marker genes was observed after 48 hours (p<0.02) and 72 hours (p<0.02). Figure 3 visualizes these gene expression changes as heatmaps (Figure 3A and 3B) and bubble plots (Figure 3C) for all genes in the plasmatocyte and lamellocyte marker gene sets.

Interestingly, a significant decrease in plasmatocyte marker gene expression can be detected before a significant increase in lamellocyte marker gene expression (Figure 3C).

To assess to what extent plasmatocyte and lamellocyte marker genes are direct targets of Ush in S2 cells we analyzed an Ush-GFP ChIP-seq dataset [17]. Indeed, 56% (24 out of 43) of plasmatocyte marker genes and 68% (34 out of 50) of lamellocyte marker genes are occupied by Ush-GFP (Supplementary figure 1D) suggesting that Ush regulates a significant proportion of these genes directly.

To verify the reciprocal regulation of plasmatocyte and lamellocyte marker genes we analyzed expression of selected genes by RT-qPCR (Figure 4).

**Fig. 4:**
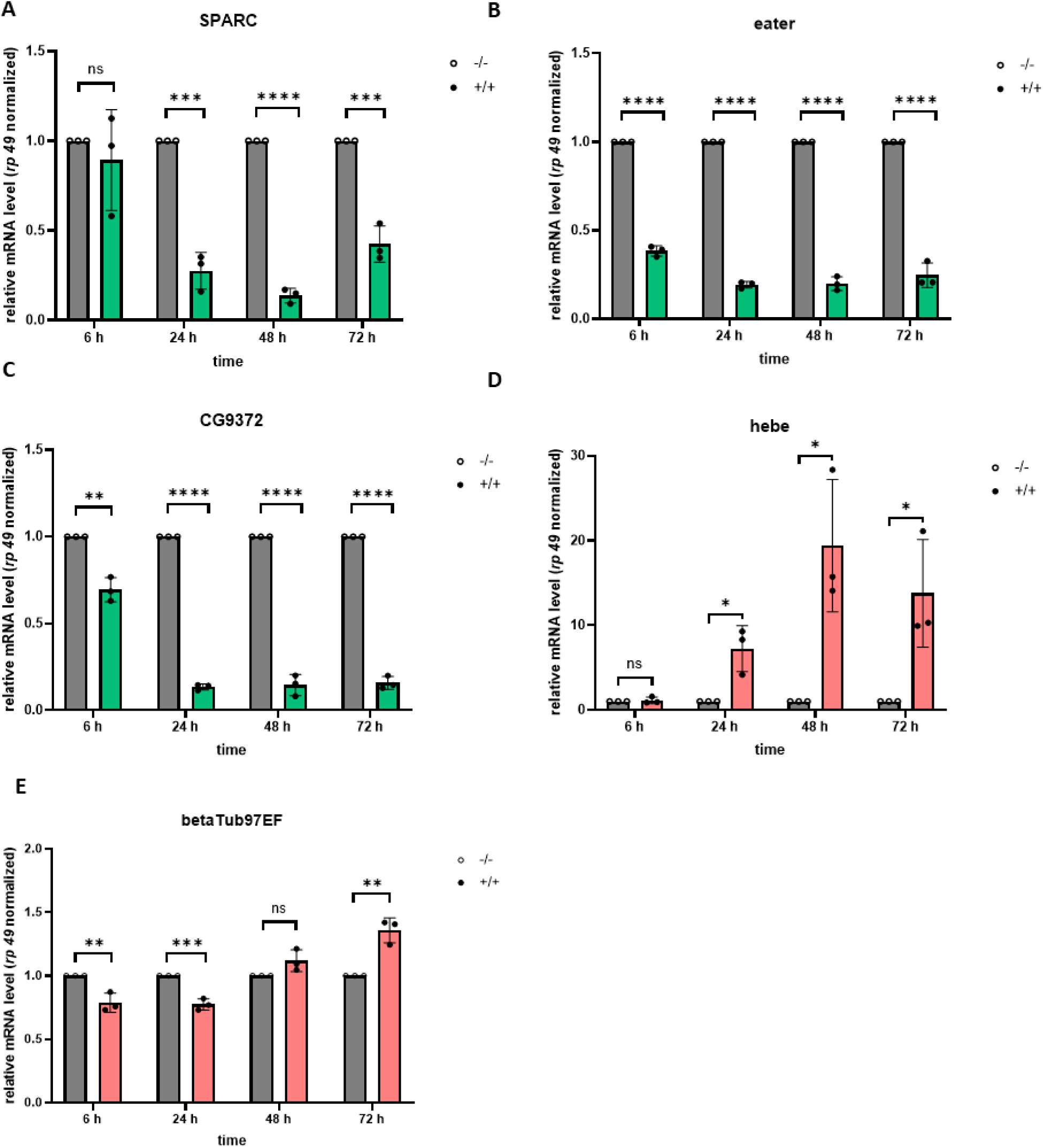
Degradation of Ush deregulates lamellocyte and plasmatocyte marker genes. RT-qPCR was used to analyse mRNA levels of *SPARC* (**A**), *eater* (**B**), *CG9372* (**C**), *hebe* (**D**) and *betaTub97EF* (**E**) in S2 Ush-GFP-D cells following induction of Ush degradation (5 µM Shield-1, 100 µM CuSO_4_) at the time points indicated below the graph. mRNA levels were normalised to *rp49* mRNA levels. mRNA levels in untreated cells were set to 1 (grey bars). mRNA levels in treated cells (green and red bars) were expressed relative to these values. Green bars represent plasmatocyte marker genes. Red bars represent lamellocyte marker genes. Individual data points and standard deviations are shown. An unpaired t-test was used for statistical analysis (ns = not significant, * p < 0.05, ** p < 0.01, *** p < 0.005).

The three plasmatocyte marker genes *SPARC*, *eater* and *CG9372* were robustly downregulated 24 hours after induction of Ush-GFP degradation (Figure 4 A-C). Indeed, *eater* and *CG9372* downregulation was detectable as early as 6 hours after induction (Figure 4 B and C). We also followed expression of three lamellocyte marker genes - *atilla* (Figure 2B), *hebe* and *betaTub97EF* (Figure 4D-E). *hebe* was upregulated 24 hours after induction and *hebe* mRNA levels further increased at later time points (Figure 4D). *atilla* expression was not significantly changed after 6 and 24 hours but increased dramatically from 48 hours onwards (Figure 2B). No increased expression of *betaTub97EF* could be detected until the 72 hour time point (Figure 4E).

Taken together, our results suggest that downregulation of plasmatocyte marker genes might generally precede the upregulation of lamellocyte marker genes. Moreover, these findings also imply a switch from a plasmatocyte-like to a more lamellocyte-like gene expression programme that is driven by a progressive decline in Ush levels.

### Ush degradation changes S2 cell morphology and adhesion

S2 cells can grow as round cells in suspension and only loosely attach to the surface of culture dishes. They express very little α-integrin [22,23], and, thus, only weakly adhere to ECM-coated substrates such as Vitronectin (Supplementary Figure 2; [24]). We used light and immunofluorescence microscopy to detect potential changes to S2 cell morphology and integrin-based adhesion upon degradation of Ush. As expected, Ush degradation resulted in a reduction of GFP signal (Figure 5A).

**Fig. 5:**
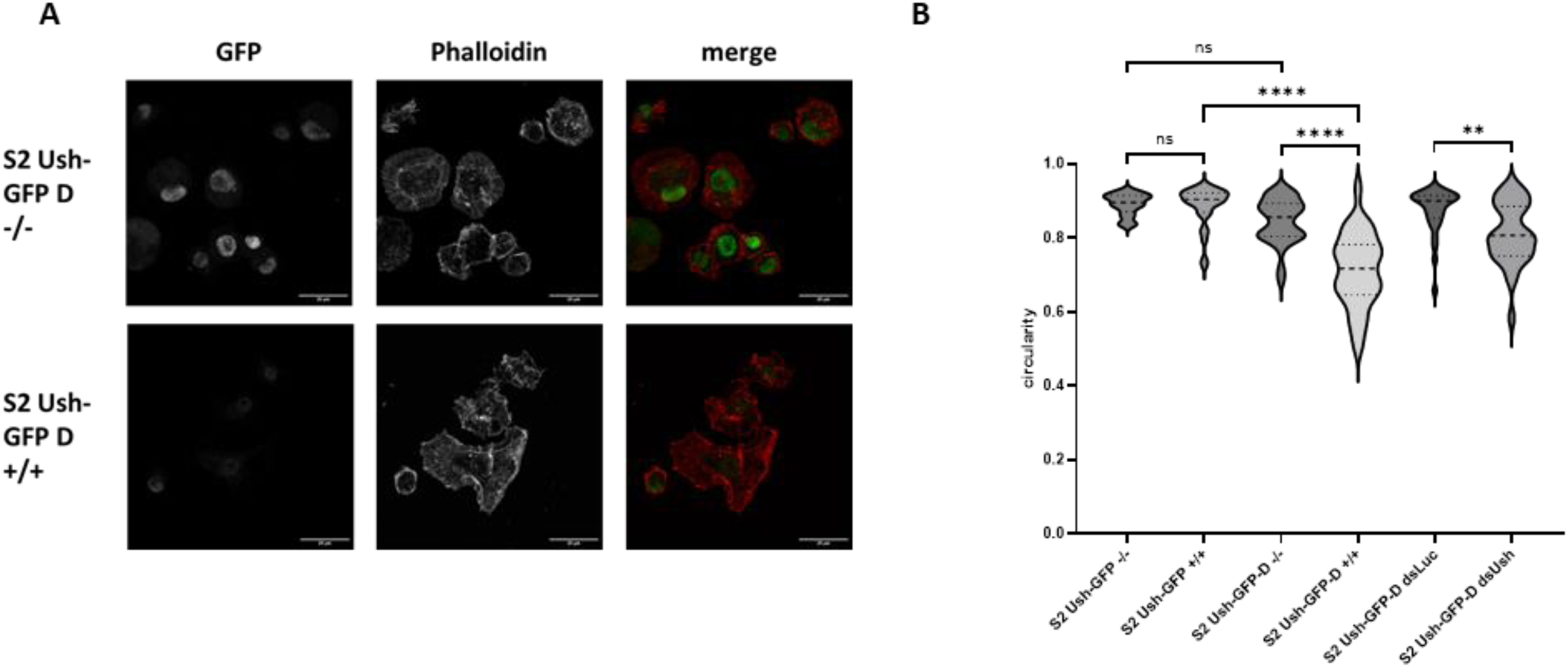
Depletion of Ush alters S2 cell morphology. **A.** S2 Ush-GFP (parental cell line) and S2 Ush-GFP-D (Degrader) cells were treated with 100 µM CuSO_4_ and 5 µM Shield-1 (+/+) for 72 h or left untreated (-/-) as indicated on the left. GFP and Phalloidin signals were acquired by fluorescence microscopy. Phalloidin stains F-actin filaments. Scale bars represent 20 µm. **B.** Circularity analysis data is represented with a violin plot. S2 Ush-GFP and S2 Ush-GFP-D cells were treated with 100 µM CuSO_4_ and 5 µM Shield-1 (+/+) for 24 h to induce Ush-GFP degradation or left untreated (-/-) as indicated below the figure. In addition, S2 Ush-GFP and S2 Ush-GFP-D cells were treated with double-stranded RNA targeting luciferase (control, dsLuc) or Ush (dsUsh) for 24 h to deplete Ush as indicated below the graph. For every condition the mean of 5 independent experiments was calculated. Circularity values were determined with ImageJ. A perfect circle has the maximum circularity value of 1. A One Way ANOVA was used for statistical analysis (ns = not significant, * p < 0.05, ** p < 0.01, *** p < 0.005, **** p < 0.001).

We also observed that many cells became more adherent and flattened out into irregular shapes (Figure 5A, Supplementary Figure 2). We used live cell imaging to visualize this morphological transition over time (Supplementary material, movies 1 and 2). To better quantify these morphological changes evoked by Ush degradation, we determined a circularity value of S2 cells under different conditions. We found no significant decrease in circularity when S2 Ush-GFP control cells were treated with Shield-1 and CuSO_4_ (Figure 5B). Likewise, there was no significant difference between the circularity of untreated S2 Ush-GFP and untreated S2 Ush-GFP-D cells. By contrast, circularity decreased dramatically when Ush degradation was induced in S2 Ush-GFP-D cells. To verify that the observed changes in cell morphology were caused by a decrease in Ush concentration we also determined circularity values following RNAi-mediated depletion of Ush in S2 Ush-GFP-D cells. Again, this resulted in a robust decrease in circularity.

The weak adhesion of S2 cells to vitronectin-coated coverslips has been attributed to a low expression of integrins [24]. Overexpression of integrins increases S2 cell attachment and spreading on vitronectin [24]. Given that Ush degradation resulted in increased spreading and adhesion of S2 cells we hypothesized that integrin expression was altered. The *Drosophila* genome encodes five integrin alpha subunits (*mew*, *if*, *scb*, *ItgaPS4* and *ItgaPS5*) and two integrin beta subunits (*mys* and *Itgbn*). Indeed, analysis of our RNA-Seq data sets revealed that all seven integrin genes of the *Drosophila* genome were deregulated upon Ush degradation: *if*, *scb*, *ItgaPS4* and *Itgbn* were all upregulated whereas *mew*, *ItgaPS5* and *mys* were downregulated (Figure 6A).

**Fig. 6:**
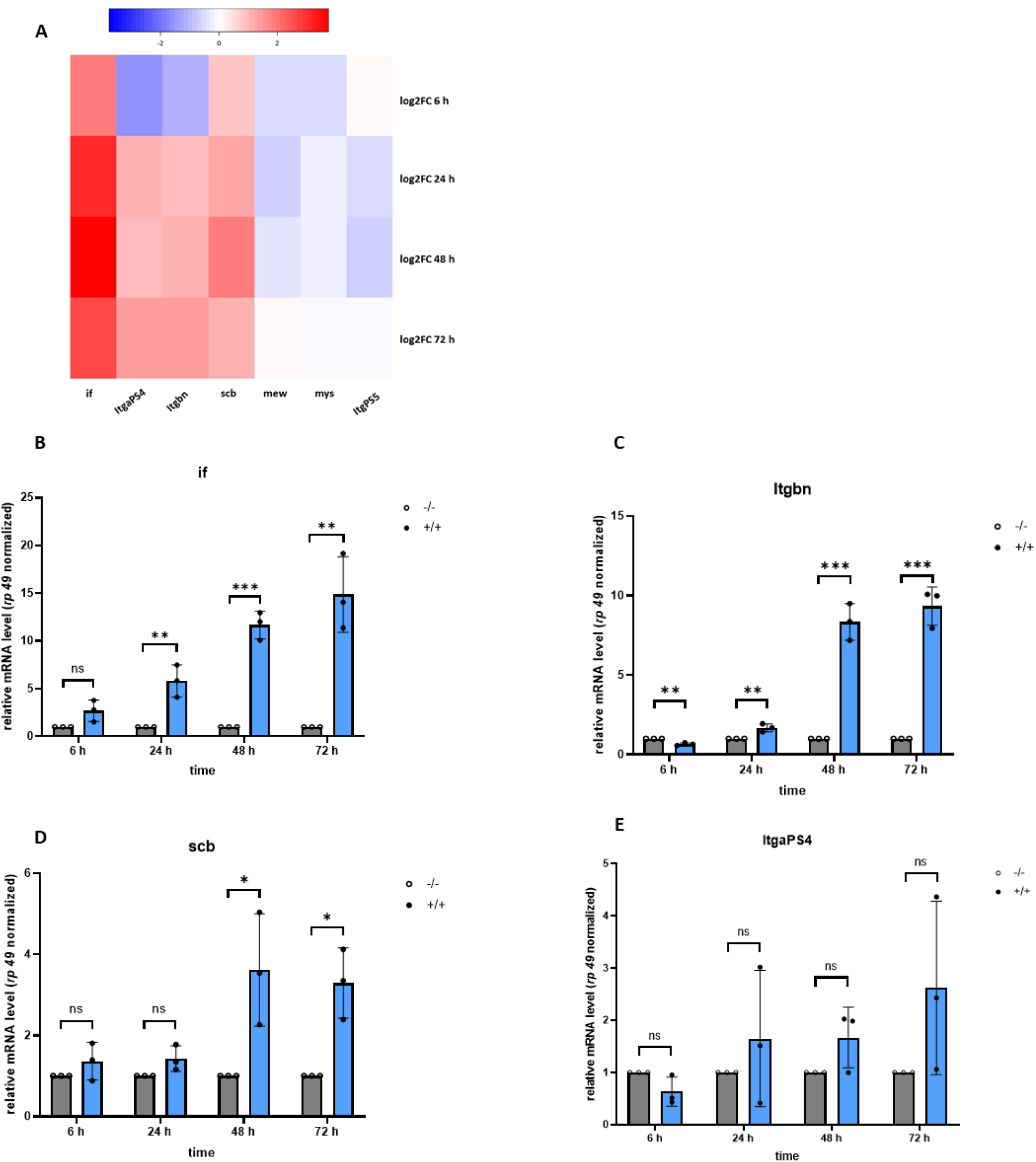
Degradation of Ush deregulates integrin expression. S2 Ush-GFP (n=3) and S2 Ush-GFP-D (n=3) cells were treated with 5 µM Shield-1 and 100 µM CuSO_4_ to induce Ush-GFP degradation or left untreated. RNA-Seq was used to determine mRNA expression differences 6, 24, 48 and 72 h after induction. Log2-fold changes of all *Drosophila* integrin genes were calculated and visualized by heat maps (A). RT-qPCR was used to analyse mRNA levels of *if* (B), *Itgbn* (C), *scb* (D) and *ItgaPS4* (E) in S2 Ush-GFP-D cells following induction of Ush degradation (5 µM Shield-1, 100 µM CuSO_4_) at the time points indicated below the graph. mRNA levels were normalized to *rp49* mRNA levels. mRNA levels in untreated cells were set to 1 (grey bars). mRNA levels in treated cells (blue bars) were expressed relative to these values. Individual data points and standard deviations are shown. An unpaired t-test was used for statistical analysis (ns = not significant, * p < 0.05, ** p < 0.01, *** p < 0.005).

To verify this finding, we monitored changes to mRNA levels of *if*, *scb*, *ItgaPS4* and *Itgbn* by RT-qPCR (Figure 6B-E). Transcription of all four genes was increased upon Ush degradation. *if* mRNA levels were modestly elevated 24 hours after induction and increased further until the 72 hour time point (Figure 6B). *Itgbn*, *scb* and *ItgaPS4* expression was not altered within the first 24 hours but rapidly increased from the 48 hour mark onwards (Figure 6B-C). Thus, these results demonstrate that transcription of integrin genes changes during Ush degradation. Integrin gene expression changes followed similar kinetics as lamellocyte marker gene expression changes (Figure 4).

We next analyzed changes in integrin localization and intensity in cells by immunofluorescence. We visualized the localization of *if* 48 hours after induction of Ush degradation in Ush-GFP-D cells grown on vitronectin-coated surfaces (Figure 7A).

**Fig. 7:**
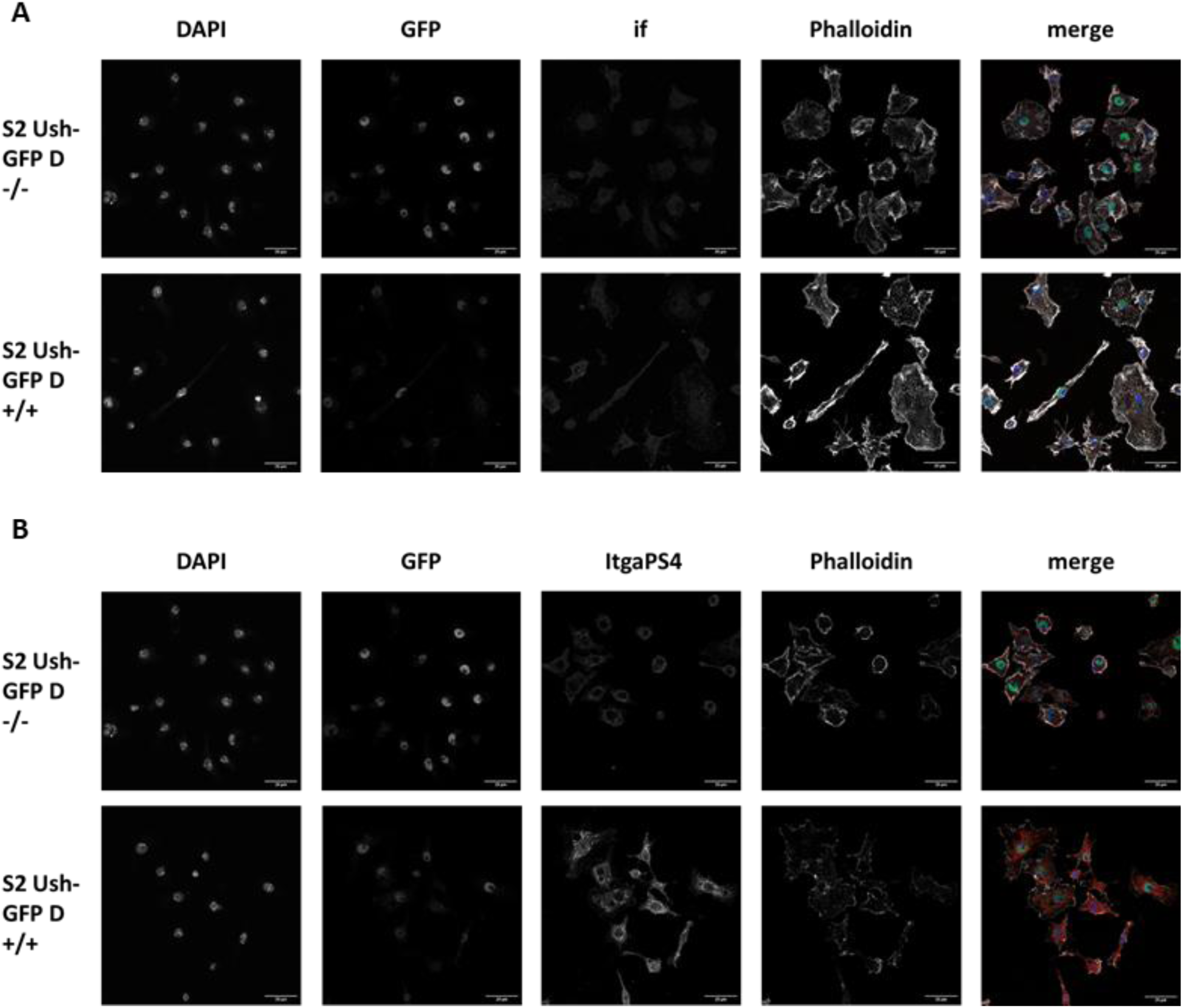
Depletion of Ush alters S2 cell adhesion. S2 Ush-GFP-D (Degrader) cells were treated with 100 µM CuSO_4_ and 5 µM Shield-1 (+/+) for 48 h or left untreated (-/-) as indicated on the left. DAPI, GFP, Phalloidin, if (**A**) and ItgaPS4 (**B**) signals were acquired by fluorescence microscopy. Scale bars represent 20 µm.

Confocal microscopy analysis revealed an overall weak punctate expression pattern of If (αPS2) and ItgαPS4 integrin in untreated S2 cells (Figure 7A, B). In contrast, after depletion of Ush cells showed increased accumulation of both α-integrins in the perinuclear region and in prominent puncta along the leading edge of cells (Figure 7A, B), characteristic for focal adhesion sites as described previously [24,25]. Thus, Ush degradation leads to changes in integrin mRNA and protein expression, resulting in an increased number of integrin-mediated focal adhesions that promotes cell spreading.

In conclusion, our analysis shows that induced degradation of Ush in S2 cells results in changes to gene expression, proliferation and matrix adhesion, all of which are reminiscent of changes occurring when plasmatocytes transdifferentiate into lamellocytes.

## Discussion

*Drosophila* larvae respond with a rapid increase of lamellocyte numbers to parasitic wasp infection. Lamellocytes are derived from two sources: from hemocyte progenitor cells in the lymph gland by regular differentiation and from fully differentiated plasmatocytes by transdifferentiation [9]. The molecular mechanisms that mediate transdifferentiation are not well understood.

Cell identity is determined by expression of genes that confer cell type-appropriate properties and by the concomitant repression of genes that define different cell types. Accordingly, plasmatocyte marker genes as defined by Hultmark and Ando are much more strongly expressed in plasmatocytes compared to other hemocyte cell types [3]. Ush degradation results in the downregulation of plasmatocyte marker genes suggesting that Ush is required to maintain their expression and, thereby, plasmatocyte-like cell identity in S2 cells. This is surprising given that prior genetic analyses have established Ush as a repressor of gene transcription that cooperates with GATA transcription factors [2]. By contrast, there is no evidence that it can also directly activate transcription. Interestingly, however, more than half of the plasmatocyte marker genes are bound by Ush suggesting that there might be mechanisms that allow Ush to directly activate transcription. Alternatively, the effects of Ush degradation on plasmatocyte marker gene expression could be mediated indirectly, for example by upregulation of repressors of plasmatocyte marker genes.

While Ush degradation decreases expression of genes that contribute to a plasmatocyte cell identity, expression of lamellocyte marker genes is activated. This is in line with previous work that has established that Ush suppresses lamellocyte differentiation by repressing lamellocyte marker genes. Mechanistically, Ush is recruited to DNA bound GATA transcription factors and cooperates with other corepressors to silence transcription [2]. Indeed, 68% of the lamellocyte marker genes analyzed in this study are bound by Ush in S2 cells.

It is noteworthy that during the degradation time course the decrease of plasmatocyte marker gene expression can be detected before the increase in lamellocyte marker gene expression (6 – 24 hours after induction versus 48 – 72 hours). It seems reasonable to assume that Ush levels diminish gradually over the 72 hour degradation period. We hypothesize that plasmatocyte marker genes require relatively high Ush concentrations to maintain their expression and are sensitive to a moderate decrease in Ush levels. By contrast, repression of lamellocyte marker genes is maintained at moderately reduced Ush concentrations. Lamellocyte marker gene derepression requires still lower Ush levels that are only achieved at later time points of the Ush degradation protocol. It is interesting to note that Ush levels do indeed progressively decrease as hemocytes differentiate *in vivo*: Hemocyte progenitors show high Ush expression, Ush levels are lower in plasmatocytes and lowest in lamellocytes [18]. This suggests that hemocyte gene expression programmes are, in part, determined by the precise concentration of Ush. Moreover, these observations suggest that during transdifferentiation the loss of plasmatocyte identity might precede the acquisition of lamellocyte identity. Induction of lamellocyte differentiation *in vivo* in response to infection with parasitic wasp eggs is a remarkably rapid process. Lamellocytes become detectable 12-24 hours after infection and their number peaks after 48 hours. The transcriptional reprogramming towards lamellocyte identity *in vivo* is faster compared to our cell culture model indicating that factors in addition to Ush are involved. Nevertheless, it is interesting to speculate that an induced degradation (or inactivation) of Ush might facilitate the rapid lamellocyte differentiation response *in vivo*.

Gene expression changes induced by Ush degradation have profound effects on S2 cell morphology: the usually round and non-adherent S2 cells attach and spread out into irregular cell shapes. Integrin expression is low in S2 cells which contributes to their weak adhesion properties [22,24]. Forced expression of integrins allows S2 cells to spread on extracellular matrices [22,26,24]. Indeed, upon Ush degradation the transcription of four of the seven *Drosophila* integrin genes (*if*, *scb*, *ItgaPS4* and *Itgbn*) are upregulated and we have demonstrated elevated protein levels for both *if* and *ItgaPS4*. These findings support the hypothesis that increased integrin expression in Ush depleted cells drives cell adhesion and promotes cell spreading. This identifies Ush as a new regulator of ECM interactions.

The ability to efficiently adhere to, spread over and encapsulate parasitic wasp eggs is a hallmark of lamellocyte function. Lamellocytes achieve this, at least in part, by expressing a unique combination of integrins. A proteomic analysis has recently identified the expression of *ItgaPS4*, *ItgaPS5* and *Itgbn* in lamellocytes [27]. It is noteworthy that two of these, *ItgaPS4* and *Itgbn*, were also upregulated in S2 cells following Ush degradation. Thus, the changes to S2 cell morphology caused by Ush degradation might mimic fundamental lamellocyte properties.

In contrast to terminally differentiated lamellocytes which no longer divide, larval plasmatocytes retain their proliferative potential. They become mitotically active in response to infections and tissue damage [9]. By contrast, degradation of Ush rapidly attenuates the proliferation of plasmatocyte-like S2 cells in concert with activating lamellocyte marker genes and increasing adhesion. We have previously shown that Ush is required for expression of genes driving the cell cycle in S2 cells including CDK1 and Cyclin B [17]. S2 cells depleted of Ush accumulate in G2 phase [17]. Genes driving the cell cycle have not been identified as plasmatocyte marker genes [3]. This is not surprising given that mitotic Cyclin/CDK complexes are expressed in all proliferating cells, not just in plasmatocytes. However, we postulate that these genes are an integral part of the plasmatocyte gene expression programme that is controlled by Ush.

It is remarkable that degradation of a single hematopoietic regulator is sufficient to change the plasmatocyte-like identity of S2 cells into a lamellocyte-like identity both at the levels of gene expression, morphology and proliferation. This underscores the importance of Ush in regulating hemocyte differentiation that has been well documented in many genetic studies [17]. It also reveals an unexpected plasticity of the S2 cell line. Interestingly, a recent in depth comparison of the transcriptomes of Kc and S2 cells has come to the conclusion that while Kc cells possess an exclusive plasmatocyte identity, S2 cells possess a certain level of transcriptional plasticity [13]. Our results suggest that S2 cells could serve as a cellular model to study transdifferentiation from plasmatocytes to lamellocytes [9]. This novel model offers unique opportunities for the future identification of the molecular mechanisms that underlie transdifferentiation.

## Materials and Methods

### Cell culture

*Drosophila melanogaster* S2 [Cas9 Ush-GFP] (abbreviation: S2 Ush-GFP) and S2 [Cas9 Ush-GFP pFFkana] clone #E3 (abbreviation: S2 Ush-GFP-D) cells were cultured in Schneider’s *Drosophila* Medium (21720024, Gibco) supplemented with 10% (v/v) fetal bovine serum (FBS; batch 0718V, Sigma-Aldrich) and 1% (v/v) Penicillin-Streptomycin (15140122, Gibco). Cell lines were grown under standard conditions at 26°C.

### Stable transfection and antibiotic selection of *Drosophila* S2 cells

S2 Ush-GFP cells were seeded in a 10 cm cell culture dish to a final concentration of 2*10^6 cells. The cells were transfected with the plasmid pMT_FKBP_L106P_Fbox_GFPbinder_kana_opt. FuGeneHD (E2311, Promega) was used as transfection reagent. The transfection was carried out according to manufacturer’s instructions. The ratio of FuGeneHD transfection reagent to DNA was 4:1. Three days post transfection the cells were transferred to cell culture medium with 800 µg/ml Geneticin (10131035, Gibco) for antibiotic selection. After 3 weeks of antibiotic selection the polyclonal cells were used for monoclonal selection. Cells were isolated by serial dilution in a 96 Well plate containing a mixture of fresh and conditioned cell culture medium. After 2 weeks monoclones were picked, analyzed by Western Blot and expanded for further use.

### RNA interference in *Drosophila* S2 cells

The MEGAscript T7 kit (AM1334, Invitrogen) was used to synthesize double-stranded RNA (dsRNA) according to manufacturer’s instructions. Briefly explained, dsRNA was generated using T7 Polymerase *in vitro* transcription from PCR amplicons obtained with T7 minimal promotor containing primers using a cDNA template from S2[Cas9] cells. 15 μg of dsRNA was added to 1.0 x 10^6 S2 cells in a total of 2.5 ml Schneider’s *Drosophila* Medium. For different cell numbers, the amount of dsRNA and medium was scaled accordingly. Cells were harvested for RNA isolation and protein extraction four days post transfection.

### Induction of GFP-Degron system

In order to stabilize the F-box protein [FKBP(L106P)-Nslmb-vhhGFP4 fusion protein] [20] and to achieve binding of the GFP-tagged protein, the small molecule inhibitor Shield-1 (HY-112210, MedChemExpress) was added to the cells with a final concentration of 5 µM. For sufficient induction of the GFP-Degron system CuSO_4_ (P023.1, Roth) is added to cells to a final concentration of 100 µM. Cells were harvested for RNA isolation and protein extraction at the indicated time points post induction.

### Cell growth assay

S2 Ush-GFP and S2 Ush-GFP-D cells were seeded in a 6-Well-cell culture plate to a final concentration of 1.5*10^6 cells. The GFP-Degron system was induced as described above. The RNAi was induced as described above. The cells were harvested at the respective time point. Prior to counting in a Neubauer counting chamber the cells were diluted 1:20 in 1 ml total volume with cell culture medium.

### Cell morphology assay

Cells were seeded and treated the same way as for cell growth assay. Pictures were taken using an Olympus CKX53 microscope extended with Olympus U-CMAD3 and Olympus UTV1X-2 camera adapters. The 20 x magnification was used. Images were acquired by an Olympus E-PL10 camera. For every condition 30 cells were measured in ImageJ with the polygon selections tool. The counting was started in the far left corner and from top to bottom until the number 30 was reached. The cells had to be in one plane. The circularity value for each cell was obtained via the ROI (region of interest) manager.

### Immunohistochemistry staining

Glass coverslips were coated with 5 µg/mL vitronectin (Sigma Aldrich) for 2h at 37°C followed by an incubation o.n. at 4°C.

After the indicated time points stimulated cells were seeded on vitronectin coated glass surface coverslips for 1-2h, fixed for 15 min with 4% paraformaldehyde (Sigma Aldrich) in PBS, permeabilized for 2-5 seconds in 0.1% Triton X-100 (Invitrogen) in PBS, blocked for 30 min in 3% BSA in PBS and subsequently stained with anti-integrin antibodies for 1h followed by secondary-antibodies (Alexa-Fluor® 568), Phalloidin-Alexa-Fluor®-647 and DAPI for 2 h.

The coverslips were then mounted on object slides using Mowiol (Roth).

Primary antibodies against the following integrins were used as follows:

anti-mys/ βPS-integrin (mouse) 1:500, (CF.6G11 from Developmental Studies Hybridoma Bank (DSHB)) anti-if (mouse) 1:100, (DSHB CF.2 C7)

anti-ItgαPS4 (rabbit) 1:100, (kind gift from C. Monod, Toulouse)

### Image acquisition and microscopy

Confocal fluorescence images were acquired with the Leica TCS SP8 with an HC PL APO CS2 63×/1.4 and HC PL APO CS2 40×/1.3 oil objective. The cells were analyzed with FIJI (ImageJ, NIH).

### Preparation of whole cell extracts

For whole cell extracts cells were resuspended and lysed in RIPA buffer (50 mM Tris pH 8.0, 150 mM NaCl, 5 mM EDTA, 1% (w/v) NP-40, 0,5% (w/v), 0,1% (w/v) SDS, 10% (v/v) glycerol; RIPA buffer is supplemented with protease inhibitors: Aprotinin 1 mg/ml, Leupeptin 1 mg/ml, Pepstatin 10 mg/ml, PMSF (Phenylmethanesulfonyl fluoride) 1 mM) for 10 min on ice while vigorously mixing the samples every 3 min. Lysates were cleared by centrifugation at 21.100 g and 4°C for 20 min. The protein content was determined using Bio-Rad Protein Assay Dye Reagent Concentrate (5000006, Biorad) according to manufacturer’s instructions. The whole cell extracts were stored at −80 °C.

### SDS-PAGE and Western Blot

Proteins were electrophoretically separated on an SDS-polyacrylamide gel (SDS-PAGE) and then transferred onto activated polyvinylidene difluoride (PVDF) membranes (T830.1, Roth) by Western Blotting in Pierce Western Blot Transfer Buffer (35040, Thermo Fisher Scientific). Membranes were saturated in blocking buffer (PBS, 0.1% (w/v) Tween-20, 5% (w/v) non-fat dry milk) for 1 h at room temperature and subsequently incubated with the respective antibody dilution in blocking buffer overnight at 4°C. After washing the membranes four times for 5 min at room temperature in washing buffer (PBS, 0.1% (w/v) Tween-20) appropriate HRP-coupled secondary antibodies (anti-mouse IgG (NA931, GE Healthcare), anti-rabbit IgG (NA934, GE Healthcare) and anti-rat IgG (31470, Thermo Fisher Scientific) were applied in blocking buffer for 2 h at room temperature. After four washing cycles for 5 min in washing buffer Western Blot signals were detected by chemiluminescence using the Immobilon Western Blot Chemiluminescence HRP substrate (WBKLS0500, Millipore). Antibodies and antisera were used in the following dilutions: GFP (1:3000; clone [3H9] from Chromotek), Tubulin beta (1:8000; clone KMX-1 from Merck Millipore), FKBP-12 (1:2000; from ThermoFisher).

### RT-qPCR and RNA-Seq

Total RNA was isolated using the peqGOLD Total RNA Kit (13-6834-02DE, Avantor/VWR) together with the peqGOLD DNase I Digestion Kit (13-1091-01DE, Avantor/VWR) and the integrity of RNA was evaluated on a 1.2% Agarose/TAE gel. For RT-qPCR cDNA was prepared from 1 μg of total RNA using the SensiFAST cDNA Synthesis Kit (BIO-65054, Bioline) and analyzed by RT-qPCR using the SensiFast SYBR Lo-ROX Kit (BIO-94050, Bioline) according to manufacturer’s instructions together with gene-specific primers (Supplementary table 1). Amplification reactions were measured in triplicates on a Stratagene Mx3000P thermocycler (Agilent Technologies) and the mean values were calculated according to the ΔΔCt method using the mRNA levels of *rp49* as a normalization reference. mRNA expression was calculated relative to non-treated samples. Error bars represent the standard deviation from three biological replicates. Statistical analysis was performed using GraphPad Prism9.

For RNA sequencing the total RNA from three independent GFP Degron system inductions and four different time points (6 h, 24 h, 48 h, 72 h) was isolated. The RNA sequencing was performed by the company Biomarker Technologies (BMK) GmbH (Münster, Germany). RNA-Seq libraries were prepared using a Hieff NGS Ultima Dual-mode mRNA Library Prep Kit (Illumina; library type unstranded). The library was inspected by Qsep-400. The Sequencing was performed on an Illumina Novaseq X platform, paired-end-read 150 bp.

### Bioinformatical analysis

The data for this study has been deposited in the European Nucleotide Archive (ENA) at EMBL-EBI under accession number PRJEB87114 (https://www.ebi.ac.uk/ena/browser/view/PRJEB87114). The accession numbers are in supplementary table 6.

Bioinformatical analysis of RNA-Seq data was carried out on usegalaxy.org. RNA-Seq data were aligned to *Drosophila* transcriptome using RNA Star (2.7.11a) [28]. Counts per gene were determined using FeatureCounts (2.0.3) [29]. Differentially expressed genes and normalized reads were determined using DeSeq2 (2.11.40.8) [30]. DeSeq2 output files were annotated using the Annotate DESeq2/DEXSeq output tables (1.1.0). Heatmaps were created using heatmap2 (3.1.3.1+galaxy0) [31]. Volcano plots (0.0.6) were created using Volcano plot [32]. The gtf-file (Drosophila_melanogaster.BDGP6.32.109_UCSC.gtf) was obtained from the Berkley *Drosophila* Genome Project (release 6, 2014-07).

GSEA (Gene Set Enrichment Analysis) [33,34] is a method determining if a predefined gene set shows statistically significant and consistent differences between two biological conditions (e.g. phenotypes “untreated” and “treated”). The “number of permutations” was set to “1000”. “Collapse/Remap to gene symbols” was set to “No_Collapse”. The “Permutation Type” was set to “gene_set”. Gene sets database see supplementary table 2. The basic and advanced fields settings were kept at default settings.

The bubble plot was created with MATLAB (Version R2021b) using the GSEA data from supplementary figure 2 A and B. The detailed MATLAB script was added to the extended data.

Supplementary figure 1E was created as follows: Fastq files of Ush-GFP ChIP and single-end reads were retrieved from Gene Expression Omnibus (ncbi.nlm.nih.gov/geo/; GSM4379724 & GSM4379725) and aligned to the *Drosophila melanogaster* genome version dm6 using Bowtie2 (2.5.3) [35]. Mapped reads were deduplicated using Samtools markdup (1.20) [36]. Peaks were called against input using MACS2 callpeak (2.2.9.1) [37] with the following settings: lower mfold: 5, upper mfold: 50, fragment size: 500 bp, minimum FDR: 0.05. Manipulation of BED files was done using bedtools (2.31.1) [38]. Peaks less than 500 bp apart were merged (MergeBED-d500). Gene positions from TSS to TES were retrieved from flybase.org and extended by 500 bp upstream of the TSS to include putative promoter regions. To identify Ush-occupied genes, Ush binding sites were intersected with extended gene positions (required overlap of 1 bp).

**Supplementary Fig. 1: Degradation of Ush deregulates lamellocyte and plasmatocyte marker genes**

Gene Set Enrichment analysis of A. plasmatocyte (PM), B. lamellocyte (LM) and C. crystal cell (CC) marker genes 6 h, 24 h, 48 h and 72 h after induction of Ush degradation in S2 Ush-GFP-D cells. A positive NES (normalized enrichment score) indicates an enrichment of the analysed gene set at the top of the ranked list. A negative NES indicates an enrichment of the analysed gene set at the bottom of the ranked list. A p-value of p < 0.02 is considered significant. D. Ush occupancy at hemocyte markers and integrin genes. A gene is considered occupied by Ush (green) if it contains an Ush ChIP-seq peak the extended gene region (gene body + 500 bp upstream of TSS).

**Supplementary Fig. 2: Depletion of Ush alters S2 cell morphology**

Brightfield pictures of S2 cells. S2 Ush-GFP (control) and S2 Ush-GFP-D cells were treated with 5 µM Shield-1 and 100 µM CuSO_4_ (+/+) or left untreated (-/-) for 24 h. S2 Ush-GFP-D cells were subjected to Ush and Luc (control) RNAi for 24 h, respectively. Representative images from the 5 replicates are shown. Images were taken with a 20x objective lens with an Olympus CKX53 microscope. Scale bars represent 20 µm.

**Movie 1: S2 Ush-GFP-D Control**

S2 Ush-GFP-D cells were left untreated. Live cell imaging was performed for 16 h. Left video showed the Brightfield channel. Right video tracked the GFP signal. Scale bar represents 20 µm.

**Movie 2: S2 Ush-GFP-D Induced**

S2 Ush-GFP-D cells were treated with 5 µM Shield-1 and 100 µM CuSO_4_. Live cell imaging was performed for 16 h. Left video showed the Brightfield channel. Right video tracked the GFP signal. Scale bar represents 20 µm.

## Supporting information

Supplementary material Ush Manuscript

## Acknowledgements

We thank Doris Wagner for technical assistance with cell culture and Tabea Trummel for assistance with MATLAB. We are grateful to Patrick Heun for donation of the Degrader expression vector and advise.

