## Supplementary material Ush Manuscript for "Targeted degradation of *Drosophila* FOG homolog U-shaped mimics macrophage transdifferentiation in S2 cells"

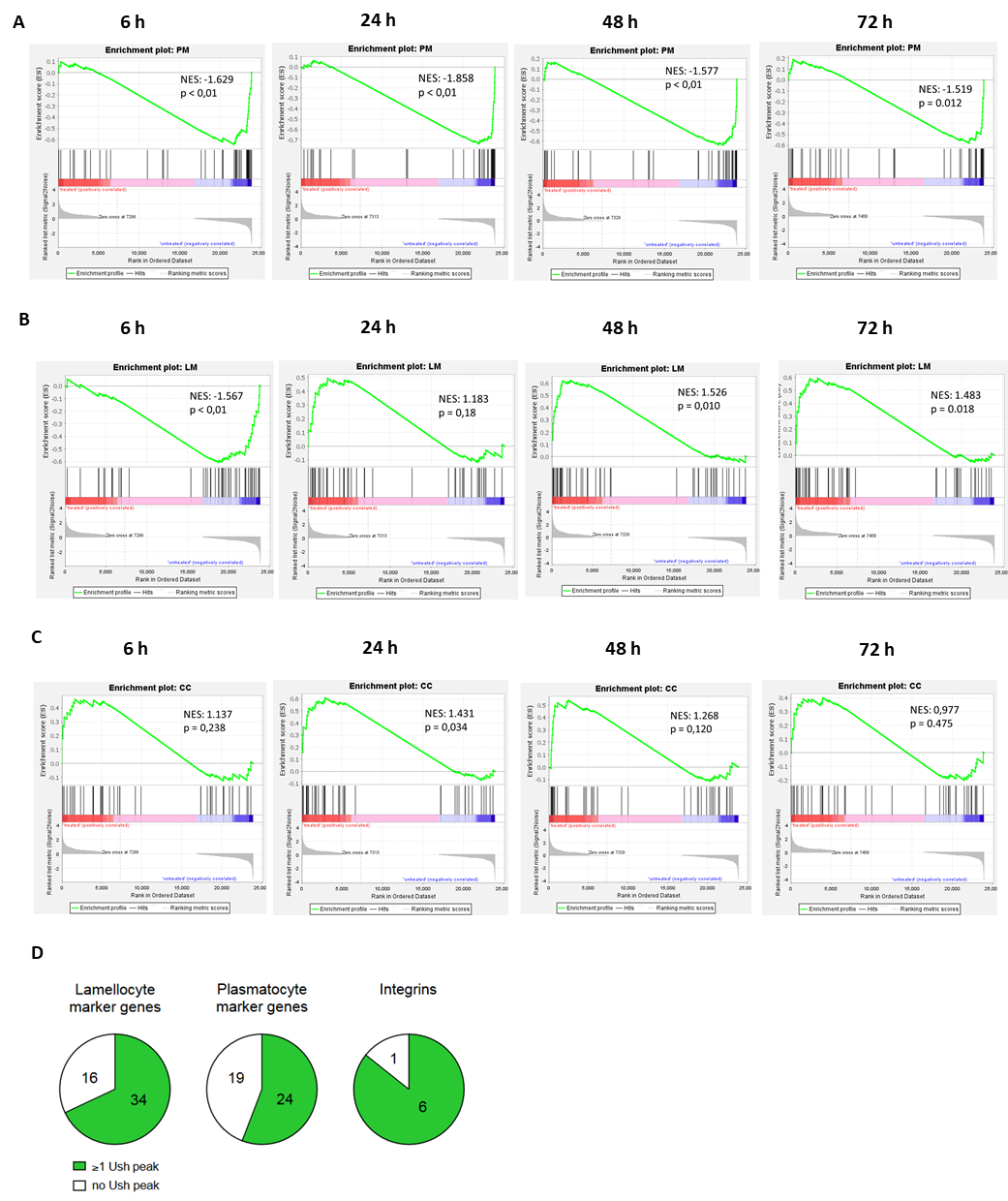

**Supplementary Fig. 1**

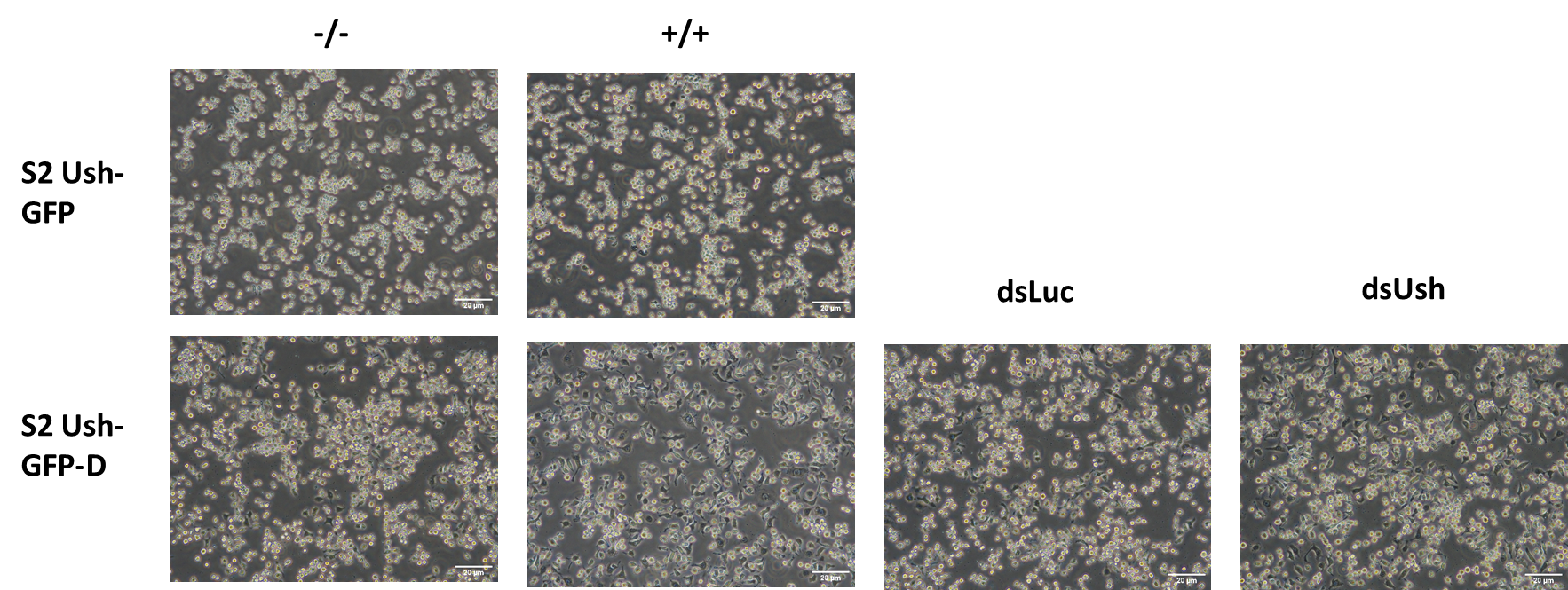

**Supplementary Fig. 2**

**Supplementary table 1: Minimal dataset for Fig. 2A-D**

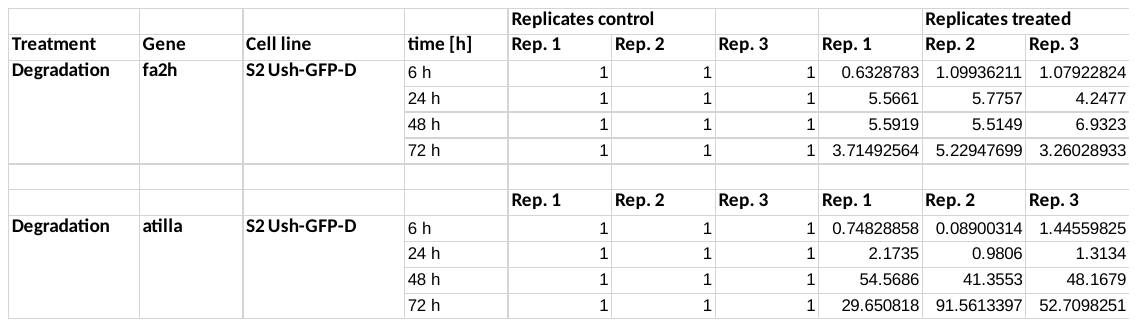

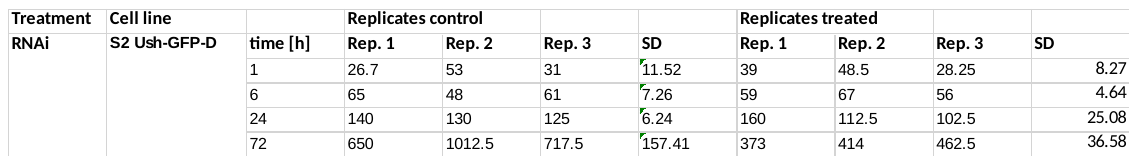

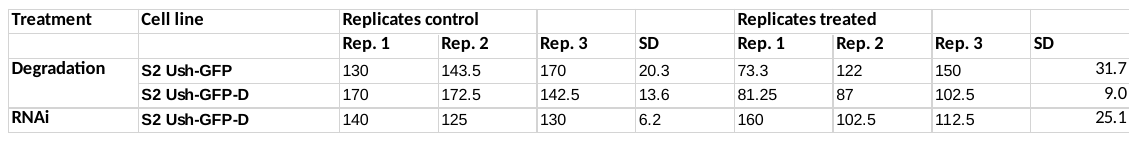

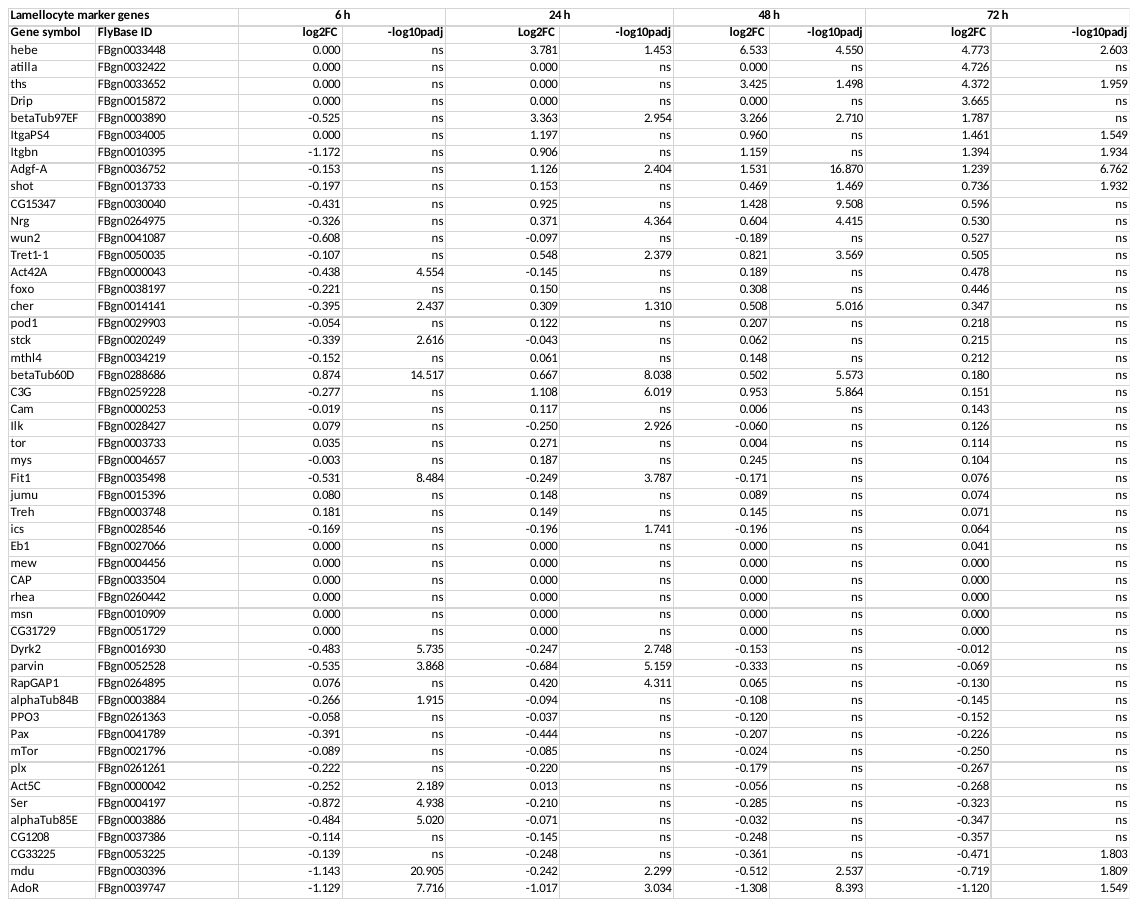
**Supplementary table 2: Lamellocyte marker gene set**

**Supplementary table 3: Plasmatocyte marker gene set**

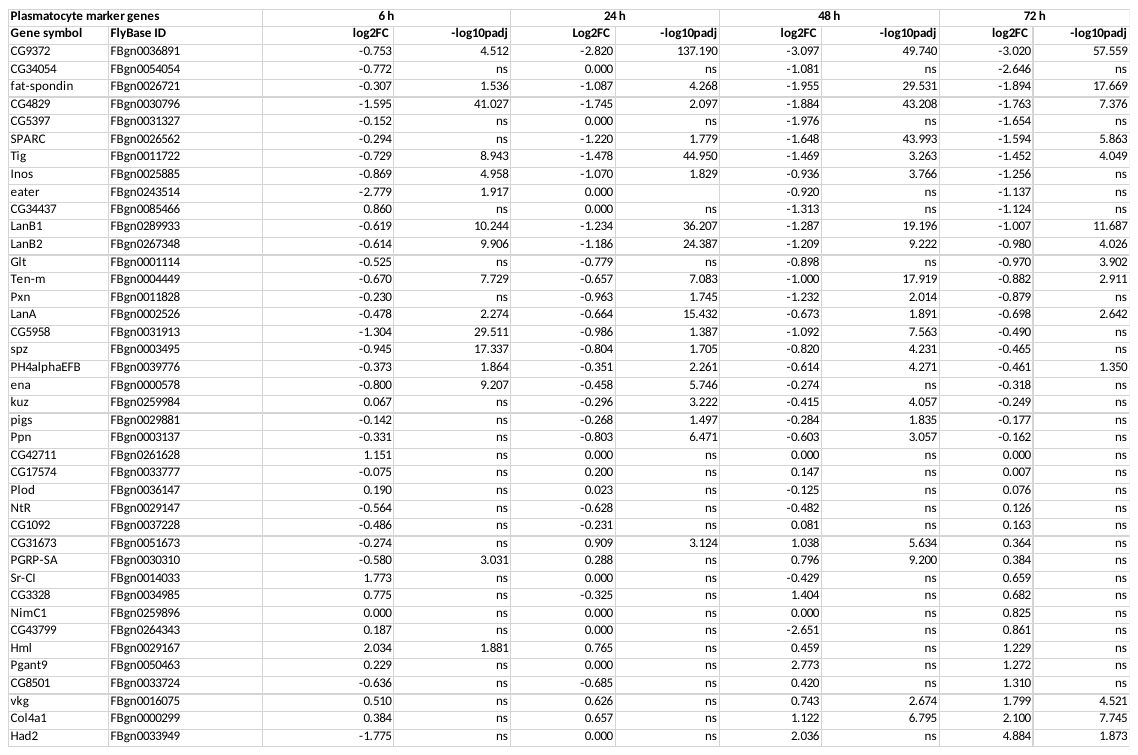

**Supplementary table 3**
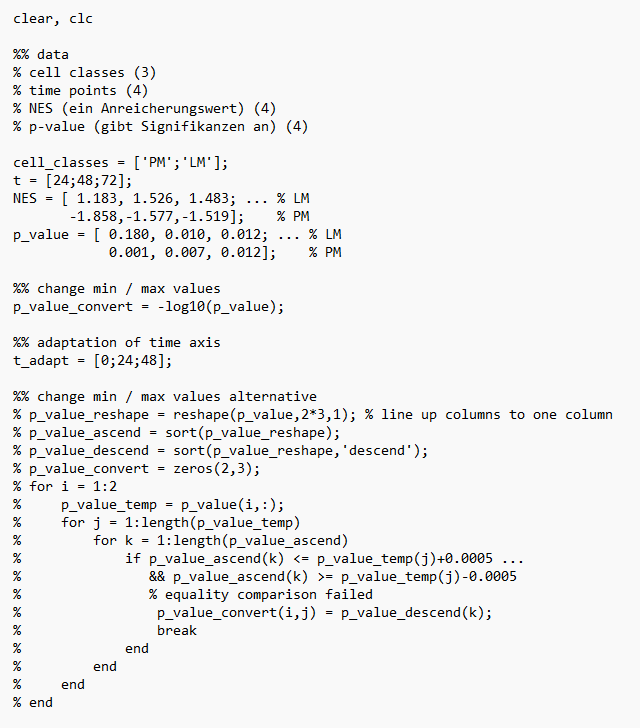
**: Minimal dataset Fig. 3C**

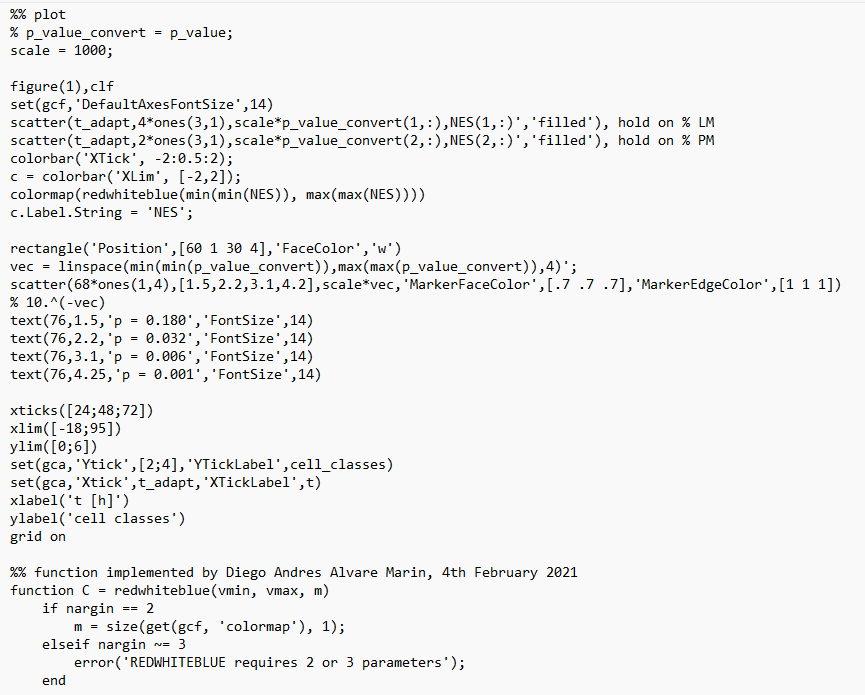

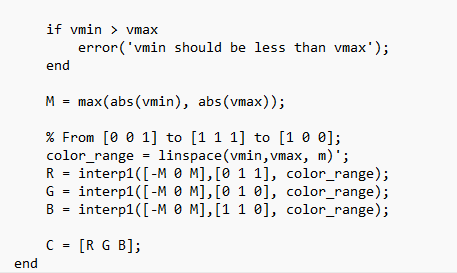

**Supplementary table 4: Minimal dataset Fig. 4**

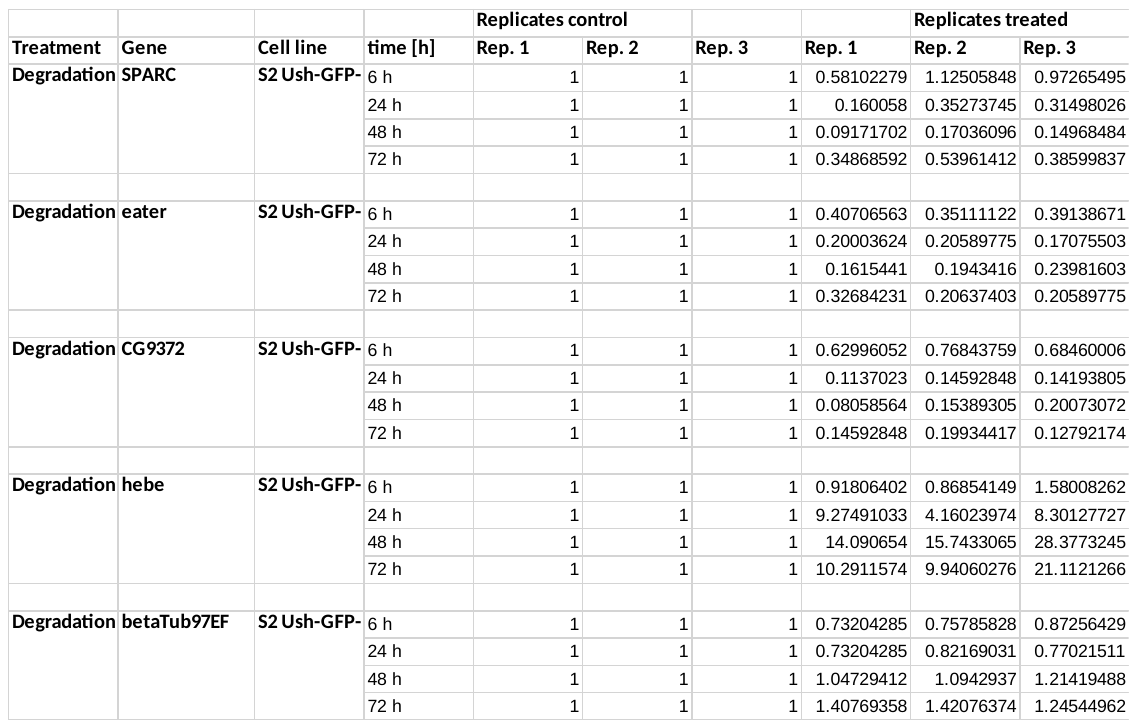

**Supplementary table 5: Minimal dataset Fig. 5 B**

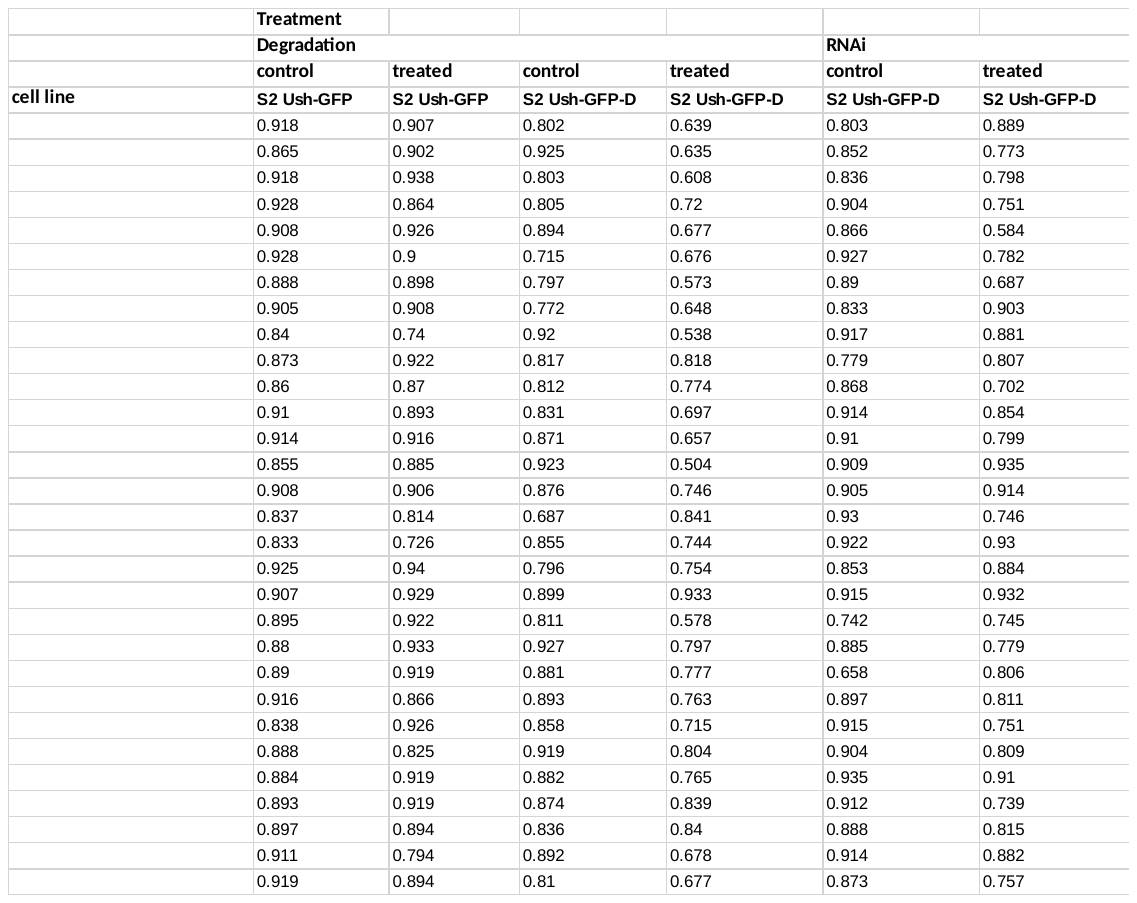

**Supplementary table 6: Integrin gene set**

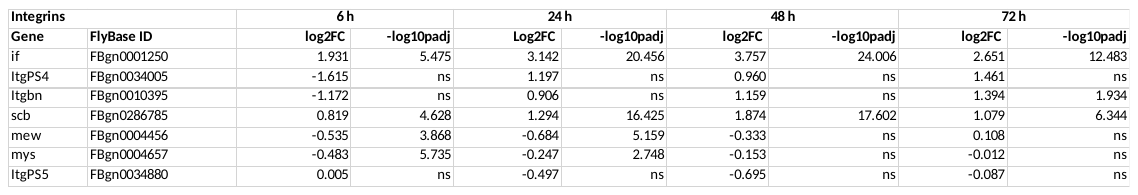

**Supplementary table 7: Minimal dataset Fig. B-E**

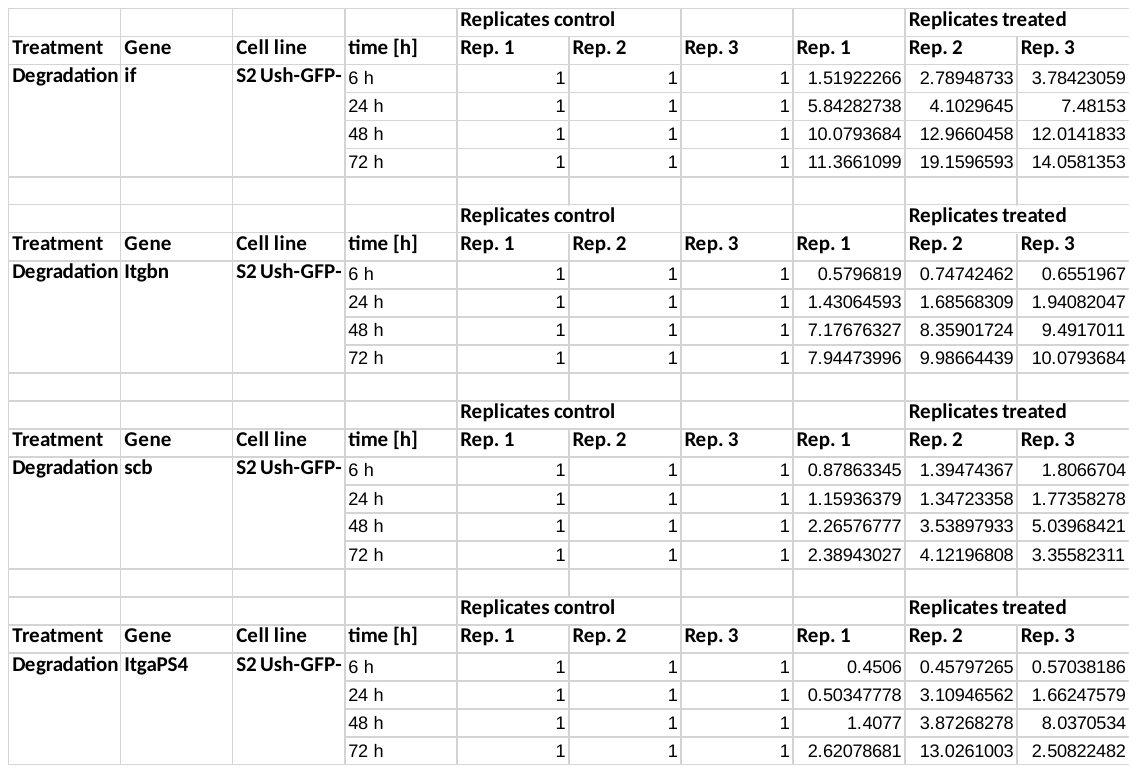

**Supplementary table 8: Gene specific primers for RT-qPCR**

| **Gene** | **Primer sequence 5‘ – 3‘** |
| --- | --- |
| fa2h-fw | GAT AGT ATG GAG CAC CTA GTG GAC |
| fa2h-rv | CCA AGG GTC AAA GAG ACG CA |
| atilla-fw | ACG GTT CAA ACC GAA AAC GAT |
| atilla-rv | TGA TGG CCG ATG CAC TGT |
| hebe-fw | GCA CAC CAA CGG ACA GAG TA |
| hebe-rv | AGA AGC TGA CCA CGG TCT TG |
| betaTub97EF-fw | GTG AAC ATG GTG CCC TTC C |
| betaTub97EF-rv | GCG CGA TAT TGC CTG GAT C |
| SPARC-fw | TCA ATC CCG TTG TCG AGG TG |
| SPARC-rv | GGA ATG CAC ACG CAT TTG GG |
| eater-fw | AGT GCG ATG ACG GTT ACA TT |
| eater-rv | GTT CCA TTT CGG CAGT TTC TTA TC |
| CG9372-fw | ACA GGT TCA ACA GCA GTA TCC |
| CG9372-rv | GAC AGT TCT CCA ACC AGC TAA A |
| if-fw | CGA TGA CTC CTA TTT GGG CTA |
| if-rv | CGA TCC TTC CCA CGA GAT TAC |
| Itgbn-fw | TCT TAG TGA CCG ATG GCT TTA TG |
| Itgbn-rv | CCG TGT ACT CAC CTG CTT TAT T |
| scb-fw | GAG TCT GCT ACT GGG TGA ATA AT |
| scb-rv | TGT TGC CGT TAT CCT CTT CC |
| ItgaPS4-fw | CGT CAA TCC AGA GGA AAG |
| ItgaPS4-rv | GTG TCG ATG TCG ATC CGA ATA A |
| rp49-fw | TGT CCT TCC AGC TTC AAG ATG ACC ATC |
| rp49-rv | CTT GGG CTT GCG CCA TTT GTG |

**Supplementary table 9: Crystal cell marker gene set**

| **Gene symbol** | **FlyBase ID** |
| --- | --- |
| CG17109 | FBgn0039051 |
| peb | FBgn0003053 |
| CG7860 | FBgn0030653 |
| Men-b | FBgn0029155 |
| CG9119 | FBgn0035189 |
| mthl10 | FBgn0035132 |
| N | FBgn0004647 |
| tna | FBgn0026160 |
| lz | FBgn0002576 |
| PPO2 | FBgn0033367 |
| PPO1 | FBgn0283437 |
| fok | FBgn0263773 |
| Galm2 | FBgn0035679 |
| MtnA | FBgn0002868 |
| CG15343 | FBgn0030029 |
| Men | FBgn0002719 |
| aay | FBgn0023129 |
| Pde1c | FBgn0264815 |
| CG5828 | FBgn0031682 |
| CG10469 | FBgn0035678 |
| Ctr1A | FBgn0062413 |
| Atox1 | FBgn0052446 |
| Gip | FBgn0011770 |
| CG10602 | FBgn0032721 |
| St3 | FBgn0265052 |
| CG5418 | FBgn0032436 |
| Fbp | FBgn0032820 |
| MtnB | FBgn0002869 |
| Fkbp59 | FBgn0029174 |
| meep | FBgn0063667 |
| path | FBgn0036007 |
| CAH2 | FBgn0027843 |
| Ald1 | FBgn0000064 |
| CG42369 | FBgn0259715 |
| CG17065 | FBgn0031099 |
| klu | FBgn0013469 |
| Naxd | FBgn0036848 |
| CG13077 | FBgn0032810 |
| CG10621 | FBgn0032726 |
| Fatp3 | FBgn0034999 |

**Supplementary table 10: European nucleotide archive accession numbers**

| **accession number** | **sample alias** | **sample title** |
| --- | --- | --- |
| ERS24000436 | D. melanogaster Unknown_BR187-001T0001 | D. melanogaster R1 6hw/o |
| ERS24000437 | D. melanogaster Unknown_BR187-001T0002 | D. melanogaster R1 6hw/ |
| ERS24000438 | D. melanogaster Unknown_BR187-001T0003 | D. melanogaster R1 24hw/o |
| ERS24000439 | D. melanogaster Unknown_BR187-001T0004 | D. melanogaster R1 24hw/ |
| ERS24000440 | D. melanogaster Unknown_BR187-001T0005 | D. melanogaster R1 48hw/o |
| ERS24000441 | D. melanogaster Unknown_BR187-001T0006 | D. melanogaster R1 48hw/ |
| ERS24000442 | D. melanogaster Unknown_BR187-001T0007 | D. melanogaster R1 72hw/o |
| ERS24000443 | D. melanogaster Unknown_BR187-001T0008 | D. melanogaster R1 72hw/ |
| ERS24000444 | D. melanogaster Unknown_BR187-001T0009 | D. melanogaster R2 6hw/o |
| ERS24000445 | D. melanogaster Unknown_BR187-001T0010 | D. melanogaster R2 6hw/ |
| ERS24000446 | D. melanogaster Unknown_BR187-001T0011 | D. melanogaster R2 24hw/o |
| ERS24000447 | D. melanogaster Unknown_BR187-001T0012 | D. melanogaster R2 24hw/ |
| ERS24000448 | D. melanogaster Unknown_BR187-001T0013 | D. melanogaster R2 48hw/o |
| ERS24000449 | D. melanogaster Unknown_BR187-001T0014 | D. melanogaster R2 48hw/ |
| ERS24000450 | D. melanogaster Unknown_BR187-001T0015 | D. melanogaster R2 72hw/o |
| ERS24000451 | D. melanogaster Unknown_BR187-001T0016 | D. melanogaster R2 72hw/ |
| ERS24000452 | D. melanogaster Unknown_BR187-001T0017 | D. melanogaster R3 6hw/o |
| ERS24000453 | D. melanogaster Unknown_BR187-001T0018 | D. melanogaster R3 6hw/ |
| ERS24000454 | D. melanogaster Unknown_BR187-001T0019 | D. melanogaster R3 24hw/o |
| ERS24000455 | D. melanogaster Unknown_BR187-001T0020 | D. melanogaster R3 24hw/ |
| ERS24000456 | D. melanogaster Unknown_BR187-001T0021 | D. melanogaster R3 48hw/o |
| ERS24000457 | D. melanogaster Unknown_BR187-001T0022 | D. melanogaster R3 48hw/ |
| ERS24000458 | D. melanogaster Unknown_BR187-001T0023 | D. melanogaster R3 72hw/o |
| ERS24000459 | D. melanogaster Unknown_BR187-001T0024 | D. melanogaster R3 72hw/ |

**Movie 1**

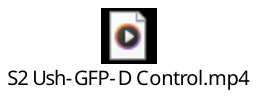

**Movie 2**

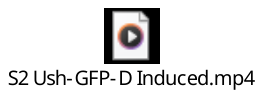
